# Sequence-Dependent Liquid Crystalline Ordering of Gapped DNA

**DOI:** 10.1101/2024.11.22.624933

**Authors:** Sineth G. Kodikara, James Gleeson, Antal Jakli, Samuel Sprunt, Hamza Balci

## Abstract

We investigate the impact of poly-adenine (poly-A) sequences on the type and stability of liquid crystalline (LC) phases formed by concentrated solutions of gapped DNA (two duplex arms bridged by a flexible single strand) using synchrotron small angle X-ray scattering and polarizing optical microscopy. While samples with mixed sequence form layered (smectic) phases, poly-A samples demonstrate a columnar phase at lower temperatures (5-35 °C), not previously observed in GDNA samples, and a smectic-B phase of exceptional stability at higher temperatures (35-65 °C). We present a model that connects formation of these LC phases with the unique characteristics of poly-A sequences, which manifest in various biological contexts, including DNA condensation and nucleosome formation.

---

Highly concentrated solutions of DNA exhibit a large variety of thermodynamic phases, including crystal and liquid crystal (i.e. mesophase) behavior ^1,2^. The rich interplay between intermolecular interactions, excluded volume considerations and possible molecular packing geometries results in an astonishing array of possibilities ^3^. Moreover, the spectrum of experimental circumstances, ranging from temperature and concentration to buffer choice, nucleic acid sequence, architecture, and end configuration, render it particularly difficult to ascertain which factor may be the most important for promoting any specific thermodynamic phase ^4^.

Short fragments of double stranded DNA (dsDNA) nominally act as rigid rods ^5,6^ and concentrated solutions of such rigid rods were predicted to form nematic liquid crystal (LC) phases if a minimum shape anisotropy (length/width≈4.7, corresponding to 28 base pair (bp) long dsDNA) is satisfied ^1,2,7^. Surprisingly, it was demonstrated that much shorter dsDNA molecules also form LC phases by stacking along their blunt ends^8–10^. Although the transition from random orientation in the isotropic phase to uniaxial order in the nematic phase results in a loss of orientational entropy, the higher translational freedom in the nematic phase and the capability to slide along the long axis of molecules more than compensate for this loss.

Above a critical DNA concentration (*c*_*DNA*_), dsDNA molecules may exhibit chiral nematic (cholesteric) and columnar LC phases^1,11^. Smectic phases, where the molecules are stacked periodically in layers along one dimension (with hexagonal in-plane positional order as in smectic-B or without it as in smectic-A), can be produced by introducing a single-stranded DNA (ssDNA) “gap” of sufficient length into an otherwise fully-paired construct^12–14^. Such “gapped” DNA (GDNA) constructs form a nematic phase over a very narrow *c*_*DNA*_ range above which smectic phases dominate^12^. When the ssDNA “gap” exceeds ∼10 nucleotide (nt) and *c*_*DNA*_ ∼260 mg/ml, a bi-layer smectic-B phase with periodicity twice the length of the GDNA duplex segment forms at or below ambient temperatures. This phase melts into a monolayer smectic-A, with periodicity comparable to a single duplex segment, at higher temperatures^13,14^.

End-to-end stacking between the blunt duplex ends and segregation of the flexible ssDNA segments between the dsDNA layers (Fig. 1A) were proposed as the primary drivers of the smectic ordering^12^. The stability of the smectic-B phase varied with the strength of the end-to-end stacking interactions, which could be modulated by varying the terminal base pairs of the duplex arms^15,16^, or by the introduction of divalent cations, specifically Mg^2+^, highlighting the significance of side-by-side interactions between neighboring duplexes^17^. These interactions are particularly important in the context of DNA condensation, and GDNA constructs provide a versatile model system to study the condensation process in a configuration where DNA molecules are naturally aligned and confined to layers.

**Figure 1.**
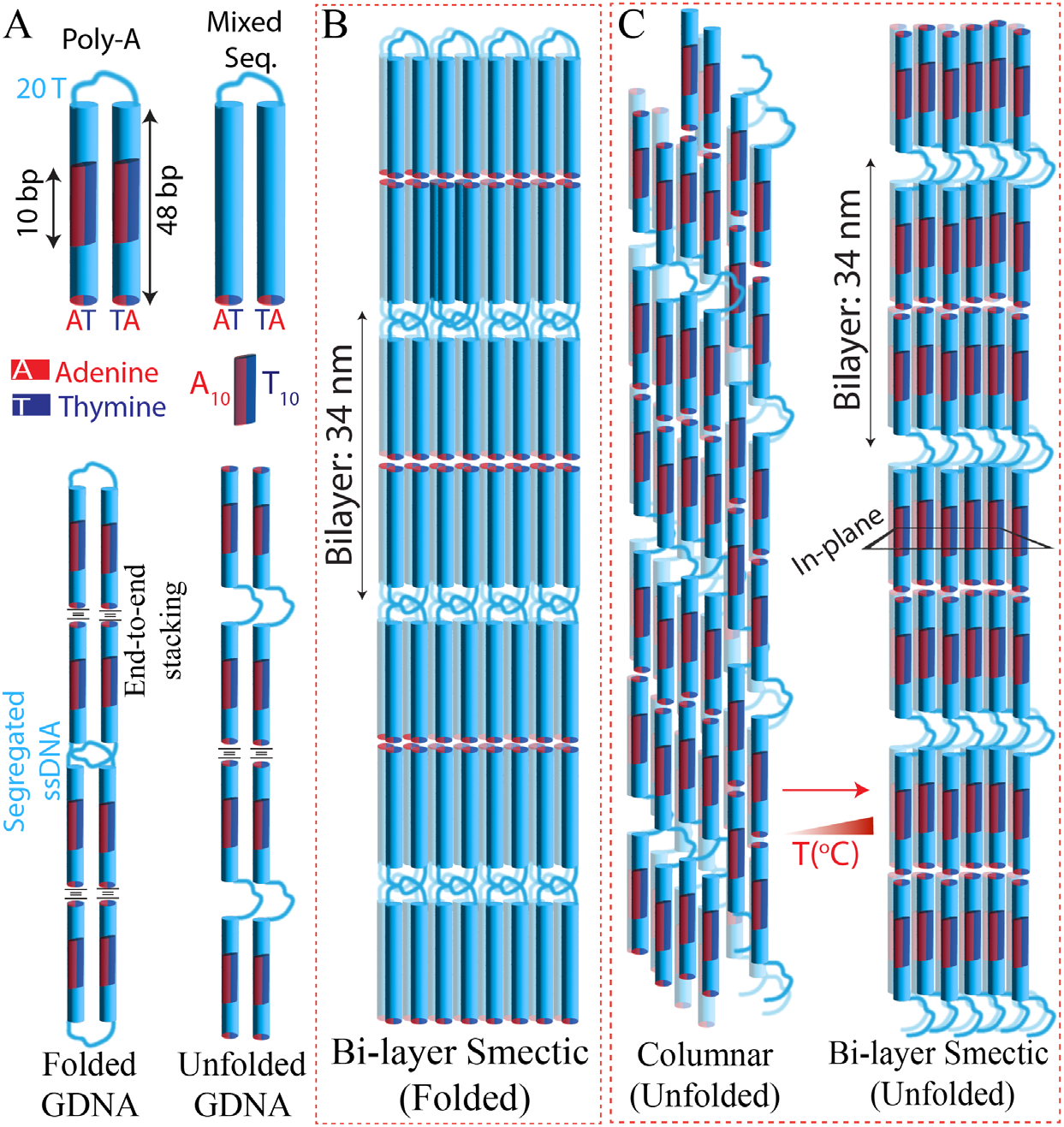
Schematics of GDNA constructs and the LC phases they form. (A) The GDNA constructs are formed by a pair of 48-bp long duplexes connected with a 20 nt long single stranded DNA linker (20T), which forms the gap region. Either a mixed sequence is used in the duplex arms or a 10 bp long region at the center is replaced with A_10_: AAAAAAAAAA (poly-A, red bar) sequence in one strand and T_10_: TTTTTTTTTT (poly-T, dark blue bar) sequence in the other. The end-to-end stacking interactions and segregation of the ssDNA gap regions are critical in the formation and stability of the bi-layer smectic phase, which can be formed by folded GDNA molecules (bottom left) or unfolded GDNA molecules (bottom right). (B) Schematics of the bi-layer smectic phase for the mixed sequence sample in the folded conformation. The layers have a high level of alignment in the direction perpendicular to the planes. (C) Schematics of the transition from the columnar phase to bi-layer smectic phase at higher temperatures for unfolded poly-A constructs. The layers in the bi-layer phase have irregularities in terms of their alignment in the direction perpendicular to the planes.

In this letter, we report the remarkable effect of ‘poly(A)-poly(T)’ enriched sequences on LC phase behavior of GNDA constructs of two 48 bp duplex segments linked by 20 nt long ssDNA (48-20T-48 design, Fig. 1A). Comparative studies were performed on two GDNA constructs, with each duplex having either a mixed sequence or a 10-bp long poly(A)-poly(T) sequence (10 A paired with 10 T, ‘poly-A’ for brevity) positioned at its center (Fig. 1A). Except for the poly-A sequences, the two constructs were identical and all measurements were performed at 150 mM NaCl. Poly-A sequences are known to facilitate the condensation process and enhance the side-by-side attraction between duplexes in the presence of cations ^18–20^. Poly-A duplexes have an intrinsic curvature, higher rigidity, and elevated bendability when such sequences periodically alternate, which impact chromatin organization and depletion of nucleosomes in poly-A sites ^21–24^. Our measurements connect these unique structural features with the impact of poly-A sequences on LC phases.

## Results

Fig. 1 and Fig. S1 show schematics of GDNA constructs with mixed and poly-A sequences forming monolayer and bi-layer smectic phases ^12,13^. Depending on whether the GDNA is folded or unfolded [27], there are multiple ways a monolayer or bilayer periodicity can be attained. These configurations are briefly discussed in Supplementary Materials (Fig. S1); however, a specific configuration is not assumed in this study.

Fig. 2A-F show temperature dependent SAXS data on mixed-sequence and poly-A samples. Diffraction from blunt end-to-end stacked duplexes of the mixed sequence constructs reveals sharp peaks at *q*_1_ ≈ 0.185 ± 0.002 nm^−1^, corresponding to a bi-layer smectic ordering with 34.0 nm layer spacing (approximately two duplex lengths), and at 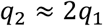, the first harmonic, at low temperature (*T* = 5 °C). Small angle peaks at similar *q* were observed for poly-A constructs only at significantly higher temperature (*T* > 35 °C). Positional correlations between duplexes within the smectic layers give rise to diffuse or sharp peaks at wide angles (wavenumber *q*_*w*_) for smectic-A or smectic-B phases, respectively. A sharp peak at *q*_*w*_ ≈ 2.15 ±0.02 nm^−1^ accompanies *q*_1_ and *q*_2_ for *T* ≲ 40 °C in the mixed sequence sample and for *T* > 35 °C in the poly-A samples, indicating smectic-B phase at these temperatures. Assuming hexagonal positional ordering of the duplexes (2.0 nm diameter) within the layers, the in-plane center-to-center spacing between the duplexes is estimated as *a = 4π/*√3*q*_*w*_ ≈ 3.37 nm (±0.03 nm).

**Figure 2.**
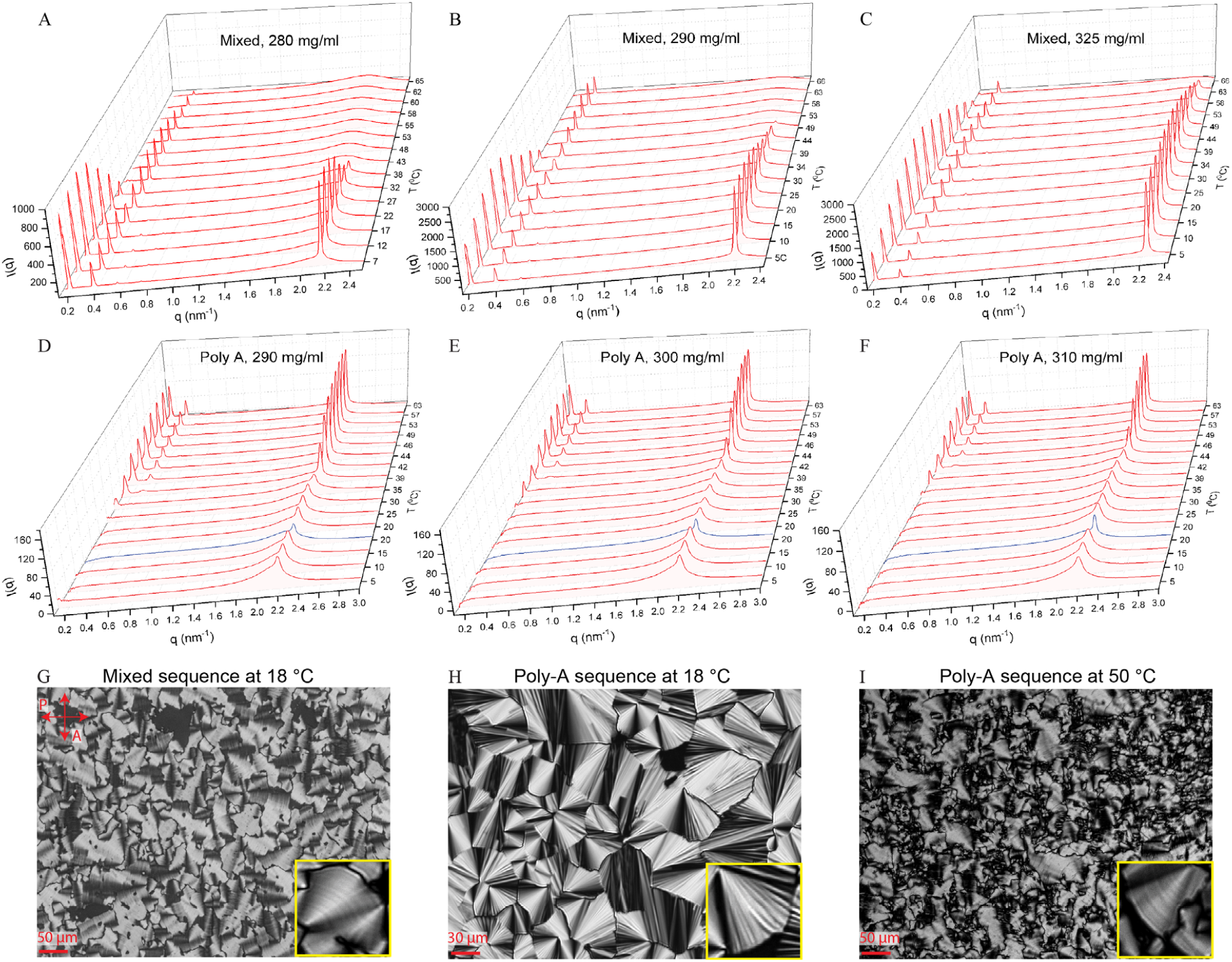
Temperature dependence (on heating from 5 °C to 65 °C) of the azimuthally-averaged SAXS intensity vs. scattering wave number *q* for the GDNA samples with mixed sequence (A-C) and poly-A sequence (D-F), in addition to POM images on these constructs (G-I). All measurements were performed at 150 mM Na^+^. The DNA concentrations were 280 mg/ml in (A); 290 mg/ml in (B); approximately 325 mg/ml in (C) for mixed sequence samples and 290 mg/ml in (D); 300 mg/ml in (E); 310 mg/ml in (F) for poly-A samples. The blue curves in (D)-(F) are the spectra after cooling the samples from 65 °C back to 20 °C. (G-I) POM images for the mixed sequence sample at 18 °C are shown in (G); poly-A sample at 18 °C in (H); and poly-A sample at 50 °C in (I). The insets show zoomed in fan textures with azimuthal striation in (G) and (I), which are consistent with a smectic phase, and stripes in radial direction in (H), which are consistent with columnar phase^2^.

In agreement with earlier observations^14,15,17^, a bi-layer smectic-B phase evolves into a monolayer smectic-A phase at higher temperatures (*T* ≈ 35 − 45 °C depending on *c*_*DNA*_) for mixed sequence samples at *c*_*DNA*_ = 280-325 mg/m (Fig. 2A-C, Fig. S2). However, similar measurements on the poly-A samples show very different patterns (Fig. 2D-F). First, the small angle peaks associated with the smectic phases are not present at low temperatures (5-35 °C) and the sharp wide-angle peak (at *q*_*w*_) is replaced with a diffuse and shallower peak for *c*_*DNA*_ = 290-310 mg/ml (Fig. 2D-F), while this feature is barely detectable at *c*_*DNA*_ = 280 mg/ml (Fig. S3). These features are consistent with a weakly ordered columnar phase in poly-A samples. At higher temperatures (T > 35 °C), a bi-layer smectic phase with long range in-plane positional correlations (sharp peaks at *q*_*1*_ and *q*_*w*_) gradually emerges and persists up to 65 °C (Fig. 2D-F). Interestingly, the columnar phase is recovered upon cooling the samples back to 20 °C (blue curves in Fig. 2D-F).

POM of poly-A GDNA solutions further support these deductions. The micrograph in Fig. 2H, taken at 18 °C on a thin layer of the poly-A solution shows the classic, radially-striped fan texture characteristic of columnar order in dense solutions of fully-paired duplexes ^1,2^. At higher temperature (T = 50 °C, Fig. 2I), the fan domains are smaller and, as highlighted in the inset, perceptively striated in the azimuthal direction. This texture is also observed in mixed sequence solutions at lower temperature (T = 18 °C) and comparable *c*_*DNA*_ (Fig. 2G) and is associated with smectic-B layering ^25^.

In Fig. 3A-B, we quantify the thermal stability of LC phases, as a proxy for the thermodynamic stability of underlying interactions, for the mixed sequence samples based on associated thermal melting temperatures (*T*_*m*_ for single-step and *T*_*m1*_ and *T*_*m2*_ for multi-step melting, listed in Table 1) as described in the Supplemental Materials [15]The results on mixed sequence samples are in-line with previous reports ^15,17^: in-plane positional order melts at slightly lower temperatures compared to bi-layer stacking and both bi-layer and in-plane orders are more stable at higher *c*_*DNA*_.

**Table 1.** Thermal melting temperatures *T*_*m*_ (obtained from single Hill function fit) and *T*_*m*1_ and *T*_*m*2_ (obtained from double Hill function fit) in units of °C. The DNA concentration is in units of mg/ml.

|  | Mixed Sequence<br>Smectic-B: Bilayer |  |  | Mixed Sequence<br>Smectic-B: In-plane |  |  | Poly-A Sequence<br>Columnar |  |  |
| --- | --- | --- | --- | --- | --- | --- | --- | --- | --- |
|  | Hill<br>Function | Double Hill<br>Function |  | Hill<br>Function | Double Hill<br>Function |  | Hill<br>Function | Double Hill<br>Function |  |
| $c_{DNA}$ | $T_m$ | $T_{m1}$ | $T_{m2}$ | $T_m$ | $T_{m1}$ | $T_{m2}$ | $T_m$ | $T_{m1}$ | $T_{m2}$ |
| 280 | 34.9±0.6 | 29.6±0.2 | 38.3±0.3 | 31.3±0.6 | 24.6±0.4 | 38.2±0.3 | N.A. | N.A. | N.A. |
| 290 | 39.2±0.6 | 32.2±0.4 | 45.6±0.4 | 37.4±0.5 | 28.4±0.5 | 45.8±0.4 | 25.9±0.6 | 18.7±0.2 | 31.2±0.4 |
| 300 | N.A. | N.A. | N.A. | N.A. | N.A. | N.A. | 31.1±0.6 | 24.0±0.6 | 36.7±0.6 |
| 310 | N.A. | N.A. | N.A. | N.A. | N.A. | N.A. | 36.4±0.5 | 32.4±0.5 | 40.2±0.6 |
| ~325 | 57.8±0.5 | 51.8±0.3 | 64.2±0.4 | 55.2±0.6 | 48.3±0.6 | 61.3±0.4 | N.A. | N.A. | N.A. |

**Figure 3.**
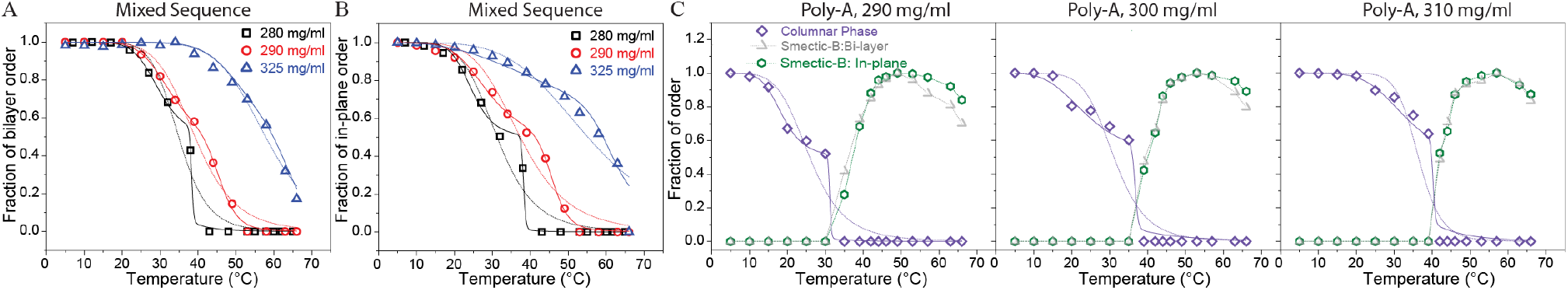
Thermal melting analysis of the SAXS data for mixed sequence (A-B) and poly-A (C) constructs. Temperature dependence of the normalized square root of SAXS intensity integrated over *q*_*1*_ and *q*_*3*_ peaks is used to quantify the fraction of the bilayer order. Similar analysis for the wide-angle *q*_*w*_ peak is used to characterize the in-plane positional order for both the smectic and columnar phases. For mixed sequence samples, temperature-dependent fraction of bi-layer order is shown in (A); and fraction of in-plane order in (B). (C) Fraction of the columnar, smectic bi-layer, and in-plane orders for poly-A samples are overlaid for the *c*_*DNA*_=290 mg/ml, 300 mg/ml and 310 mg/ml samples. The dashed and solid lines overlaid on columnar data represent single and double-Hill function fit, respectively. The sharp kinks at double-Hill functions are at positions where data points are not present, which could not be avoided due to limited temperature resolution. The dashed-lines overlaid on bi-layer and in-plane orders of smectic-B phase in poly-A constructs are guides to the eye.

For poly-A constructs (Fig. 3C), the wide-angle peak that is characteristic of the columnar phase is relatively weak and is located on top of a diffuse shoulder, which forms a non-uniform background, at T=5-35 °C. We modeled this background using a Gaussian which was subsequently subtracted to quantify the area under the superposed, sharper peak. On the other hand, the smectic phase is not present at low temperatures and appears to stabilize in the 35-45 °C range, with SAXS peak heights reaching maximum values in the 50-55 °C range and diminishing slightly (by 10-20%) at 65 °C. Higher temperatures were not tested due to denaturation of the duplexes at ∼80 °C. Therefore, it was not possible to reliably determine the corresponding *T*_*m*_ for the poly-A samples. The vanishing of the wide-angle feature characterizing the columnar phase and the emergence of the peaks characterizing the smectic-B phase coincide at similar temperatures (35-45 °C, Fig. 3C), implying a competition between these phases.

## Discussion

Our data demonstrates a picture that is consistent with a temperature dependent transition from a columnar to a bi-layer smectic-B phase for the poly-A constructs. The columnar phase was not reported in previous studies on gapped DNA constructs with mixed sequence^12–15,17^, which highlights the impact of the specific structural features of poly-A sequences on the formation and stability of LC phases^18–20,26,27^. The higher propensity of poly-A enriched duplexes to condense implies stronger effective attractive interactions between the duplexes^18^. The attractive interactions between the central poly-A region of a duplex and the mixed sequences (above or below the central poly-A) of the neighboring duplex might be stable enough at low temperatures to trap the system in a local free-energy minimum that prevents alignment of central regions and strictly planar side-to-side arrangement of the duplexes. This would frustrate the formation of ordered layer structures and frustrate the smectic phase. At elevated temperatures these relatively weak side-by-side interactions might break, allowing the system to rearrange such that the central poly-A regions are aligned. This would enable a smectic layer structure to be established (accompanied by segregation of the single stranded segments) as the global free energy minimum state. Fig. 1C schematically illustrates this transition for the unfolded configuration. Upon cooling the sample from 65 °C to 20 °C, the sample reverts back to the columnar phase (blue curves in Fig. 2D-F), suggesting the global rearrangement of the molecules is reversible. This might be due to DNA molecules occasionally exploring out of plane conformations at all temperatures but being trapped in them only at low temperatures, as described above.

A relevant observation in this context is the significant difference in amplitude of the X-ray diffraction from the bilayer stacking between the poly-A and mixed sequence samples (*q*_*1*_ amplitude ∼1000 a.u. for mixed vs. ∼100 a.u. for poly-A). The peak amplitude is an indicator of mass density contrast (Δρ) along the direction normal to the layer plane: well aligned duplexes and uniformly segregated flexible ssDNA regions would result in larger amplitude peaks due to higher Δρ, i.e., repeating pattern of high-ρ in the duplex regions followed by a low-ρ in the flexible regions. Therefore, our data suggests the duplexes are well-aligned and the gap regions well-segregated in mixed sequence samples while these are less uniform in poly-A samples. This is consistent with the duplexes of poly-A samples occasionally being trapped at non-optimal positions (e.g., duplexes invade the otherwise predominantly flexible linker region), which might frustrate the layering and result in the columnar phase at low temperatures. This model of competing columnar and smectic phases would not apply for mixed sequence samples if the side-by-side attractive interactions that trap the molecules out of plane positions are not strong enough.

The bi-layer smectic-B phase persists to ≥65 °C in the poly-A samples despite AT terminal base pairs (which produce weaker end-to-end attraction than AT-GC or GC-GC terminations) and the absence of multivalent cations, which enhance the stability of the smectic-B phase. It is quite remarkable that, at comparable *c*_*DNA*_, the thermal stability of the smectic phase in our poly-A samples surpasses even the optimal conditions for the mixed sequence system (GC-GC terminal base pairs and ∼30 mM Mg^2+^)^15,17^. A recent all-atom molecular dynamics study might provide some insights into this surprising result^19^. This study proposed a robust continuous cation distribution in the major groove of a DNA construct that includes poly(A)-poly(T) sequences in contrast to a patched 3D cation distribution observed for a mixed sequence construct. The cations (specifically Mg^2+^) located in the major groove of the poly(A)-poly(T) construct were in register with the negatively charged phosphate backbone of the neighboring duplex and demonstrated shallower binding to the major groove. Such shallow binding and orientational coupling facilitated attractive interactions with the negatively charged phosphate backbone of the neighboring duplexes, resulting in a higher propensity for DNA condensation. Even though Mg^2+^ was not included in our measurements and the computational work only reported Na^+^/Mg^2+^ mixtures, similar attractive pockets (“sticky patches”) along the duplex arms could trap the neighboring DNA molecules in positions that prevent uniform layering at low temperatures. At higher temperatures, breaking of these weak bonds might allow alignment of the poly-A segments approximately symmetrically (although with imperfect segregation of duplexes and gap regions as suggested by the smaller peak amplitude of *q*_*1*_), resulting in the smectic phase. When in approximate alignment, the stronger attractive interactions between the poly-A segments and the backbone of neighboring DNA molecules would act as a new mechanism that stabilizes the smectic-B phase and give rise to the elevated stabilities we observe.

Our comparative studies on the poly-A and mixed sequence constructs demonstrate that the underlying interactions between neighboring gapped DNA constructs can be tuned in a sequence-dependent manner. The introduction of short poly-A sequences produces a profound impact on the LC phase behavior, including the replacement of a higher order smectic (smectic-B) phase with a columnar order in the low (5-35 °C) temperature range and, surprisingly, the stabilization of smectic-B order at substantially higher temperatures. The remarkable stability of the smectic-B phase in the poly-A constructs -- to temperatures approaching the denaturing temperature of the duplexes -- highlights the significance of sequence composition and potentially of charge distributions along grooves of DNA molecules in tuning the helix-helix interactions. Our study provides a striking new example of the rich mesophase behavior of gapped DNA constructs, which continue to expand our understanding of DNA-based condensed phases.

## Acknowledgements

This study was supported by the National Science Foundation under Grant No. DMR-1904167. SAXS/WAXS measurements were performed on the CMS beamline (11-BM) at the National Synchrotron Light Source II, a U.S. Department of Energy (DOE) Office of Science User Facility operated for the DOE Office of Science by Brookhaven National Laboratory under Contract No. DE-SC0012704. The authors are particularly grateful to R. Li, F. Yang, and M. Fukuto for their assistance in these measurements.

