## Supplementary Materials for "Sequence-Dependent Liquid Crystalline Ordering of Gapped DNA"

Sineth G. Kodikara[a], James Gleeson [a], Antal Jakli [a,b,], Samuel Sprunt[a,b,*], and Hamza Balci[a,*]
[a] Department of Physics, Kent State University, Kent, OH 44242, USA;
[b] Advanced Materials and Liquid Crystals Institute, Kent State University, Kent, OH 44242.

### Materials And Methods

**Gapped DNA (GDNA) Synthesis:** DNA oligomers (O1, O2, and O3) were purchased from Genescript (Piscataway, NJ) in polyacrylamide gel electrophoresis (PAGE) purified form. The sequences of O1, O2, and O3 for both the mixed and poly-A constructs are given in Table S1. The poly-A constructs were created by replacing the mixed sequences in 20$^{th}$-29$^{th}$ nt and 88$^{th}$-97$^{th}$ nt of O1 (approximately centers of the two duplex arms) with consecutive A's while the bases in the corresponding segments of O2 and O3 were replaced with consecutive T's.

GDNA synthesis and sample loading into thin-walled borosilicate capillaries were carried out as described previously (12). The GDNA constructs (48-20T-48) consist of two symmetric 48-base pair duplexes connected by a 20 nt long single-stranded DNA gap of consecutive thymine bases (Fig.1A). To create the GDNA constructs, a long strand (O1, 116 nt) is annealed with two short strands (O2 and O3, each 48 nt) which are complementary to either side of O1 except the 20T sequence in the middle. After annealing, the GDNA solutions are passed through a 50 kDa membrane filter (Amicon Ultra from Millipore) to remove incomplete constructs and diluted such that they contain ~30 mM NaCl by adding distilled and deionized water. This solution is then concentrated by centrifuging it for 12 min at 12,000 rpm (GDNA concentration increases while NaCl concentration is kept at ~30 mM). The GDNA concentration is measured with a NanodropOne Spectrometer (Thermofisher Scientific) and is typically in the 80-100 mg/ml range at this step. This solution is then added to quartz capillaries (2 mm inner diameter) with an open end. Water is slowly evaporated which results in increasing GDNA and NaCl concentration. The NaCl concentration is adjusted such that ~150 mM NaCl will be reached when the desired DNA concentration is achieved. The only exception to this procedure is the mixed sequence sample with 325 mg/ml concentration, which was inadvertently concentrated to this level due to imperfect sealing. As described in Supplementary Information (Fig. S1), 325 mg/ml is an estimated $c_{DNA}$ and should be considered as a lower bound. Despite the uncertainty in $c_{DNA}$, this sample was included in the manuscript to illustrate the persistence of smectic phase patterns (and the absence of the columnar phase) for mixed sequence samples even at significantly higher $c_{DNA}$ compared to those studied for the poly-A constructs (280-310 mg/ml).

**SAXS Measurements:** SAXS measurements were carried out on beamline 11-BM at the National Synchrotron Light Source II. The incident X-ray energy was 17 keV, and the incident beam size at the sample was 0.2 × 0.2 mm. The typical acquisition time for SAXS patterns was 5 s, which did not result in visible X-ray damage to samples. A commercial hot/cold stage with Kapton film windows was used to regulate the sample temperature between 5-65 °C, which is well below the thermal melting temperature (~81 °C at the relevant DNA concentration) of the 48-bp duplex arms. We also recorded background scattering from a capillary containing pure buffer solution, which was subtracted from the data taken on the GDNA samples during processing of the SAXS patterns. A capillary filled with silver behenate powder was used to calibrate the scattering wave number ($q$) in the plane of the detector.

**Polarizing optical microscopy:** A droplet (~0.5 μl) of the concentrated GDNA solution (~100 mg/ml) was applied to a clean microscope slide and covered with a Deckgläser cover glass, without the use of an external spacer. The resulting film between the slides is estimated to be only a few micrometers thick. The optical texture of the sandwiched sample was observed through the microscope eyepiece as the water gradually evaporated from the open edges of the film at 18 °C in a dry laboratory atmosphere. Initially, the concentrated samples exhibited a featureless dark texture (isotropic). Over time, as evaporation occurred at the edges, the samples transitioned from isotropic to cholesteric, and further into smectic/columnar phases, characterized by bright fan-shaped textures. Upon the formation of a thick smectic/columnar rim, the sample was sealed along its edge with a thin ring of mineral oil. Regions where the mineral oil infiltrated the sample appeared as featureless islands, distinctly separated from the aqueous GDNA solution by a well-defined boundary. After sealing, the GDNA concentration within the film gradually became more uniform between the edges and the center, although cholesteric and smectic/columnar domains typically coexisted. Once the desired texture was observed, images were captured using a Nikon camera attached to the microscope. To confirm the phase transformation of the poly-A sample, the sample was placed in an Instec TS62 hot/cold stage mounted on the microscope, and images were captured at 50 °C.

| **Construct** | **Strand** | **Sequence (5′ to 3′)** | **Length(nt)** |
|---|---|---|---|
| Mixed Sequence | O1 | ACAGATGCACATATCGAGGTGGACATCACTTACGCTGAGTACTT CGAATTTTTTTTTTTTTTTTTTTTAAGCTTCATGAGTCGCATTCAC TACAGGTGGAGCTATACACGTAGACA | 116 |
| | O2 | TTCGAAGTACTCAGCGTAAGTGATGTCCACCTCGATATGTGCAT CTGT | 48 |
| | O3 | TGTCTACGTGTATAGCTCCACCTGTAGTGAATGCGACTCATGAA GCTT | 48 |
| Poly-A Sequence | O1 | ACAGATGCACATATCGAGGTTTTTTTTTTTTACGCTGAGTACTTC GAATTTTTTTTTTTTTTTTTTTTAAGCTTCATGAGTCGCATTTTTTT TTTTTGGAGCTATACACGTAGACA | 116 |
| | O2 | TTCGAAGTACTCAGCGTAAAAAAAAAAAACCTCGATATGTGCAT CTGT | 48 |
| | O3 | TGTCTACGTGTATAGCTCCAAAAAAAAAAAATGCGACTCATGAA GCTT | 48 |

**Table S1.** Sequences of the DNA oligos used to create the GDNA constructs.

**Thermal Melting Analysis:** Thermal melting analysis is performed as described in more detail previously (14). This analysis relies on the thermal evolution of the area under the relevant SAXS peaks, which are the first (fundamental) and third order peaks for the bi-layer order and the sharp wide-angle peak for the in-layer order. The cumulative area under the first and third order peaks were used to characterize the bi-layer phase. The square root of the area scales with the magnitude of the density modulation or order parameter that characterizes the degree of positional order in each case. Each peak was fit to a Gaussian function in $q$ using Origin software, and the total area under the Gaussian was then calculated (described in Supporting

Information Fig. S1). The square root of the resulting total integrated intensity under the relevant peaks is normalized with respect to the relevant maximum value and then plotted as a function of temperature in Fig. 3. The results were fit to single and double Hill functions of the form,

$f(T) = 1 - \frac{T^n}{T_m^n + T^n}$ and $f(T) = 1 - \frac{1}{2}\left(\frac{T^n}{T_{m1}^n + T^n} + \frac{T^K}{T_{m2}^K + T^K}\right)$ , where $f(T)$ is the average fraction of "bound" duplexes, $T_m$, $T_{m1}$, and $T_{m2}$ are characteristic melting temperatures, and $n$ and $k$ are positive exponents. Table 1 lists these values for all constructs we investigated. In certain cases, the double Hill function produces an artificial kink in a region where the data is sparse. With limited synchrotron beam time, it was not feasible to perform measurements at higher temperature resolution.

**Additional Schematics**

Fig. 1C shows a schematic of the transition of poly-A construct from columnar to bi-layer smectic phase for the unfolded GDNA conformation. In Fig. S1-A, we demonstrate a similar transition for the folded GDNA conformation of poly-A construct. Compared to the bi-layer smectic phase formed by the mixed sequence samples (Fig. 1B), the poly-A constructs are illustrated to possess irregularities in alignment of the duplexes in the out of plane direction. Supplementary Figure S1-B shows a schematic of the monolayer phase for the poly-A samples.

The remarkable phase transition from columnar to smectic-B raises questions about the structural characteristics of the bi-layer smectic-B phase formed by poly-A constructs, which could adopt either unfolded (Fig. 1C) or folded (Fig. S1A) conformations [27]. Based on computational simulations [11], earlier results demonstrating the prevalence of smectic phases in solutions of mixed sequence constructs have generally been modeled in terms of the folded configuration. However, the thermodynamic favoring of a columnar over smectic phase at low temperatures is more difficult to envision with folded constructs. In particular, end-to-end stacking of such constructs would tend to "cap" the column length while also frustrating the freedom of columns to slide next to each other (Fig. S1). On the other hand, longer columns could be readily formed by end-to-end stacking of the unfolded molecules (Fig. 1C, left). The substitution of columnar for smectic order at low temperature therefore suggests that poly-A sequences suppress the folded GDNA configuration.

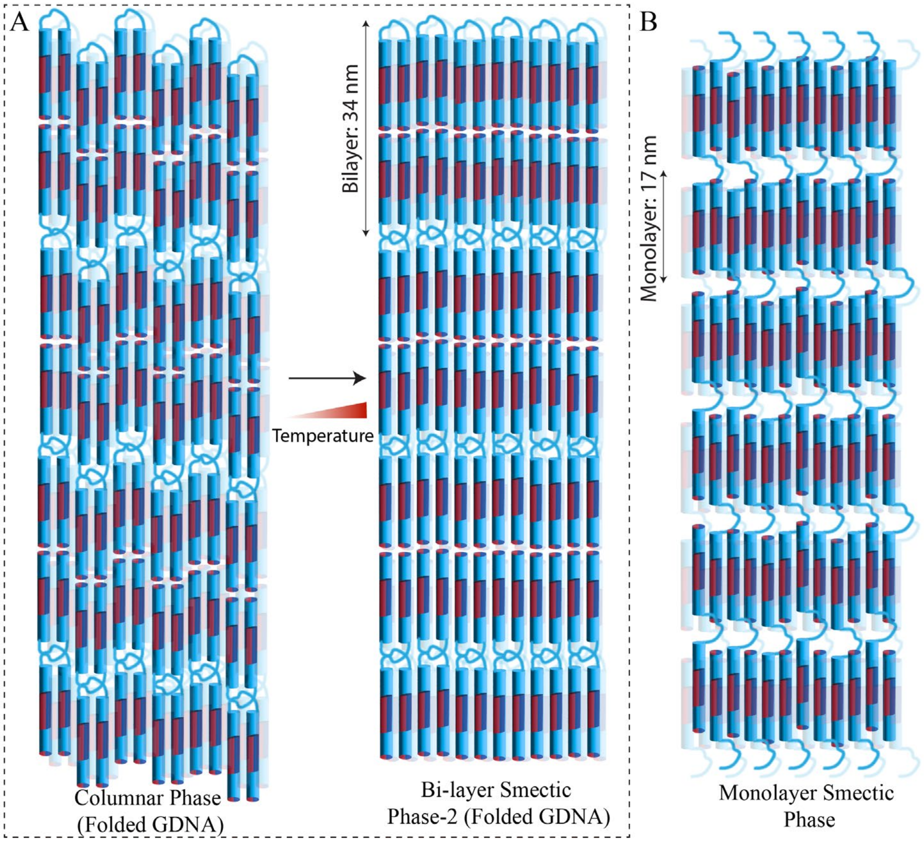


**Figure S1**. (A) Schematics of the transition from columnar to bi-layer smectic phase for the folded GDNA conformation of poly-A construct. (B) Schematics of the monolayer smectic phase for the poly-A construct.

### Estimating the Concentration of the Sample with Imperfect Seal

The constructs with a mixed sequence demonstrate a bilayer smectic-B phase at lower temperatures, followed by a monolayer smectic-A phase at higher temperatures, which eventually melts into an isotropic phase at even higher temperatures. This pattern appears to be preserved in mixed sequence samples over a broad DNA concentration ($c_{DNA}$) range as demonstrated with the $c_{DNA}$=325 mg/ml sample. This sample was inadvertently concentrated beyond the target $c_{DNA}$=300 mg/ml due to imperfect sealing of the capillary, which was realized during the measurements at the synchrotron so the concentration could not be adjusted. Due to the mentioned irregularity, it was not possible to determine DNA concentration for this sample using our usual methods. Instead, the concentration was estimated by assuming a linear relationship between the thermal melting point of the bi-layer phase and $c_{DNA}$. Using the $c_{DNA}$ and $T_m$ values of the mixed sequence construct at three lower $c_{DNA}$ values (260-265 mg/ml from an earlier study (14), and 280 and 290 mg/ml from the current study), we estimated the concentration of the current sample to be $c_{DNA}$=325 mg/ml. Supplementary Figure S2 shows this fit. Due to the presence of only three data points, the fitting errors were very large and would estimate ±40 mg/ml uncertainty in $c_{DNA}$. This is clearly an overestimate of the errors since $T_m$=57.8±0.5 °C for the sample with unknown concentration is much higher than the $T_m$=39.2±0.6 °C for the sample with $c_{DNA}$=290 mg/ml, so the DNA concentration should be significantly higher than 290 mg/ml. To illustrate, raising $c_{DNA}$ from 280 mg/ml to 290 mg/ml increased the $T_m$ by 4.3 °C (Table 1). At such a rate, $c_{DNA}$ would need to be 323 mg/ml when $T_m$=57.8±0.5 °C. Clearly, assuming a linear relationship between $T_m$ and $c_{DNA}$ may not be justified over such an extended range since we expect the $T_m$ to saturate at very high $c_{DNA}$ concentration. In that case, cDNA of the sample in question would be estimated to be higher than 325 mg/ml. Therefore, we believe the concentration of this sample is in 325-350 mg/ml range, but we quote the lower bound in the manuscript. Despite these issues with estimating $c_{DNA}$, this sample was included in the manuscript as it demonstrates the prevalence of smectic phase (and lack of transition to a columnar phase) for mixed sequence samples at $c_{DNA}$ significantly higher than those studied for poly-A constructs (280-310 mg/ml).

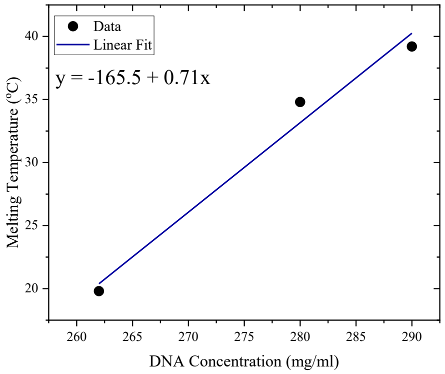


**Figure S2**. Melting temperature vs. DNA concentration for mixed sequence samples in the current and a previous study. Blue line is a linear fit to the data.

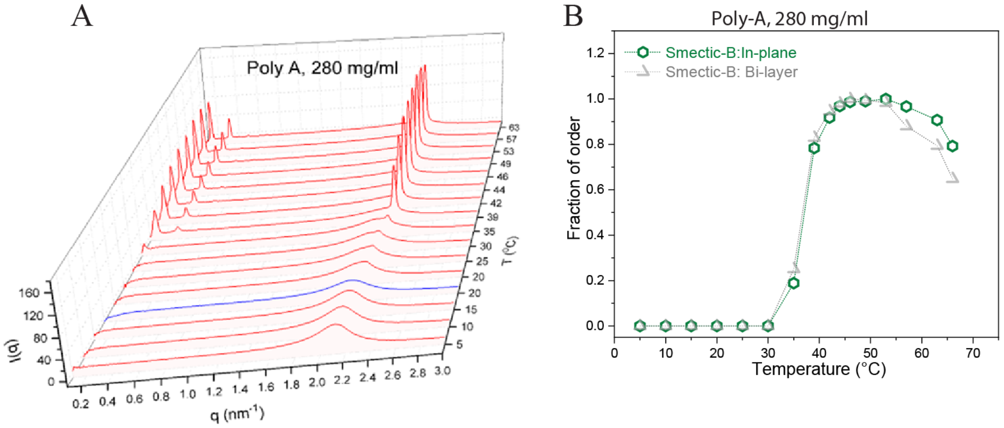


**Figure S3**. (A) Temperature dependence (on heating from 5 °C to 65 °C) of the azimuthally-averaged SAXS intensity vs. scattering wave number $q$ for the poly-A construct at $c_{DNA}$=280 mg/ml. The blue curve is the spectrum after cooling the sample from 65 °C back to 20 °C. (B) Fraction of smectic bi-layer and in-plane orders for poly-A sample at $c_{DNA}$=280 mg/ml. The dashed-lines overlaid on bi-layer and in-plane orders of smectic-B phase are guides to the eye.
